# Orientation of Temporal Interference for Non-Invasive Deep Brain Stimulation in Epilepsy

**DOI:** 10.1101/2020.07.07.191692

**Authors:** Florian Missey, Evgeniia Rusina, Emma Acerbo, Boris Botzanowski, Romain Carron, Agnès Trébuchon, Fabrice Bartolomei, Viktor Jirsa, Adam Williamson

**Affiliations:** Institute de Neurosciences des Systèmes (INS), INSERM, UMR_1106, Aix-Marseille Université, Marseille, France; Department of Functional and Stereotactic Neurosurgery, Aix-Marseille Université, Timone University Hospital, Marseille, France; Laboratory of Organic Electronics, Campus Norrköping, Linköping University, Norrköping, Sweden

**Keywords:** electrical stimulation, cortex, hippocampus, seizures, epilepsy, temporal interference

## Abstract

In patients with focal drug-resistant epilepsy, electrical stimulation from intracranial electrodes is frequently used for the localization of seizure onset zones and related pathological networks. The ability of electrically stimulated tissue to generate beta and gamma range oscillations, called rapid-discharges, is a frequent indication of an epileptogenic zone. However, a limit of intracranial stimulation is the fixed physical location and number of implanted electrodes, leaving numerous clinically and functionally relevant brain regions unexplored. Here, we demonstrate an alternative technique relying exclusively on nonpenetrating surface electrodes, namely an orientation-tunable form of temporally-interfering (TI) electric fields to target the CA3 of the mouse hippocampus which focally evokes seizure-like events (SLEs) having the characteristic frequencies of rapid-discharges, but without the necessity of the implanted electrodes. The orientation of the topical electrodes with respect to the orientation of the hippocampus is demonstrated to strongly control the threshold for evoking SLEs. Additionally, we demonstrate the use of square waves as an alternative to sine waves for TI stimulation. An orientation-dependent analysis of classic implanted electrodes to evoke SLEs in the hippocampus is subsequently utilized to support the results of the minimally-invasive temporally-interfering fields. The principles of orientation-tunable TI stimulation seen here can be generally applicable in a wide range of other excitable tissues and brain regions, overcoming several limitations of fixed electrodes which penetrate tissue.

## Introduction

During presurgical evaluation, patients suffering from focal drug-resistant epilepsy often require invasive recordings using stereo-electroencephalography (SEEG), involving the implantation of numerous electrodes in different brain regions for the electrophysiological monitoring of seizure onset and the subsequent localization of an epileptogenic zone (EZ)^1,2^. Electrical stimulation from the intracranial electrodes is often necessary to help define an EZ, with electrophysiological discharges and seizures triggered by different frequencies of stimulation^3,4^. In general, pathological discharges from the EZ are characterized by several biomarkers, primarily the generation of high-frequency oscillations in the beta/gamma range, classically referred to as rapid-discharges^5,6,7^. Although currently performed with invasive SEEG, less-invasive methods capable of evoking such discharges and seizures during the localization of EZ tissue would be highly interesting as positive surgical outcomes are well-correlated with the removal of tissue regions which generate such high-frequency oscillations^8,9,10^.

In the work presented here, we show the method of orientation-tunable Temporal Interference (ot-TI) for evoking seizure-like events (SLEs) having the characteristic frequencies of rapid-discharges. The method presents several clear advantages over classical brain stimulation and potential future applications in presurgical evaluation and treatment of human epilepsy. Our method of ot-TI can explore brain tissue, including sensitive regions previously unavailable for direct implantation, in a minimally-invasive way using electrodes on the cortex surface. Points of focal stimulation at significant distances below the cortex surface are created by envelopes of interacting electric fields applied by the surface electrodes to evoke SLEs in the hippocampus. The orientation of the electric field, as defined by the orientation of the surface electrodes, determines the effectiveness and the threshold of stimulation necessary to evoke the SLE.

Temporal interference stimulation was recently introduced in 2017^11^. The concept of TI stimulation exploits physiological properties of neurons, namely that the neuronal membranes filter electrical signals of frequencies more than 1 kHz, limiting depolarization and stimulation properties above these frequencies. In this work, we show that a crucial part of TI stimulation is the electrode configuration – not simply to move the spot of focal stimulation – but more importantly to move the orientation of electric field of the TI with respect to the orientation of the underlying structure to be stimulated. For classic TI, two electric fields are used with each field having a slightly different frequency, *f_1_* and *f_2_* = *f_1_* + Δ*f*, where f1 is selected above the threshold to evoke electrophysiological activity, for example 1200 Hz, and Δ*f* is selected at a frequency typically used to evoke activity, for example 50 Hz. The two electric fields of frequency 1200 Hz and 1250 Hz create a low-frequency envelope of 50 Hz. Classically, electrode references have been placed in the chest of animals. However, this allows field lines to pass relatively arbitrarily through subcortical structures such as the hippocampus. As we show, based on the configuration of the electrode pairs it is possible to evoke dramatically different activity exploiting the orientation of the field lines with respect to the subcortical anatomy. In Figure 1A and B, we utilize local references and orient electrode pairs with respect to axon tracts, the Schafer collaterals, of the CA field.

**Figure. 1:**
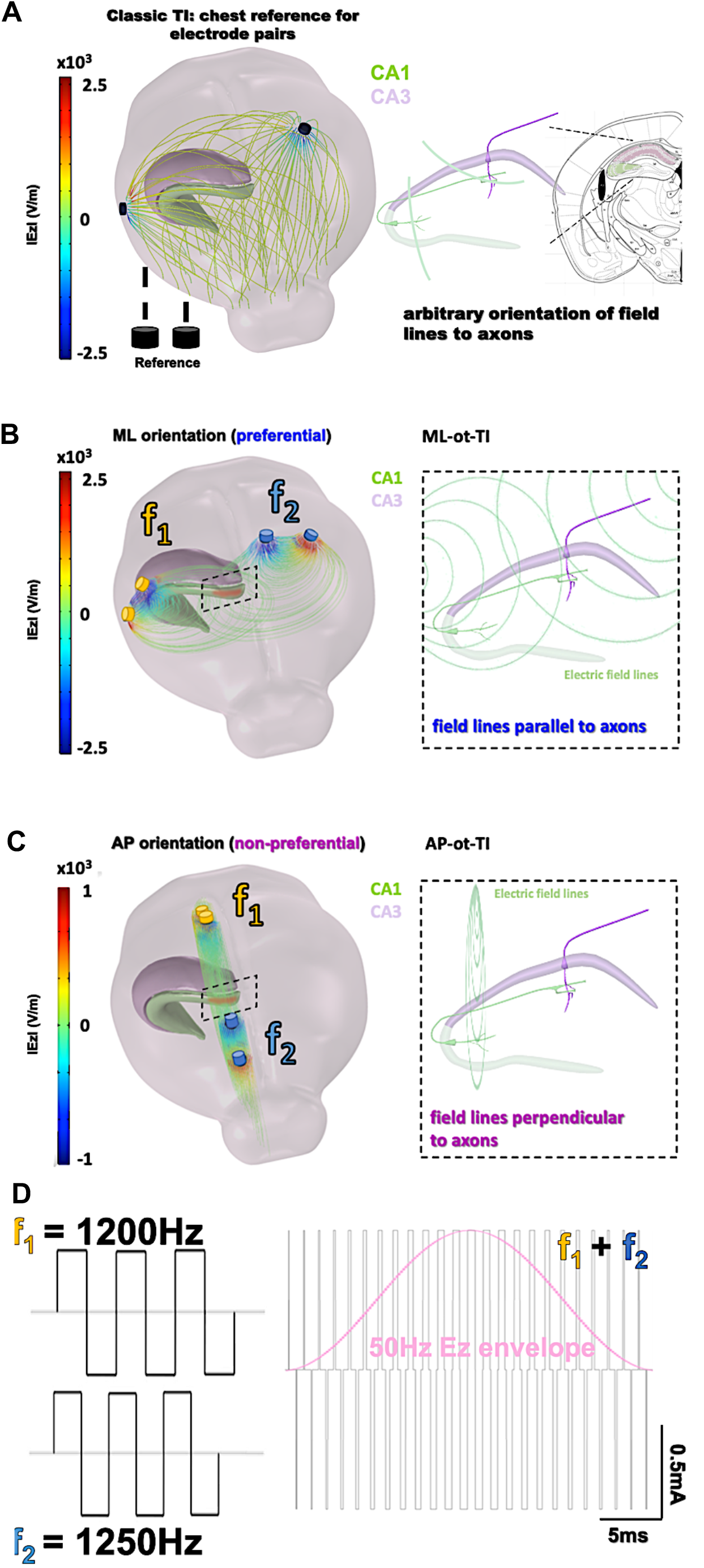
Orientation-tunable Temporally Interfering Electric Fields (ot-TI). **(A)** Classic TI, using electrode pairs with references in the chest, a reliable method to non-invasively stimulate. However, field lines are arbitrary with respect to sub-cortical structures. **(B)** ot-TI, using electrode pairs with local references for controlled-orientation of electric field lines. Here, two perpendicular arrangements of ot-TI are shown, ML with field lines parallel to the alignment of axons (inset in A) in the hippocampus from CA3 to CA1 and **(C)** AP with field lines perpendicular to the same axons (inset in B) **(D)** Classic TI stimulation is performed with sine waves, however envelopes can also be created with other wave forms, for example square waves as seen here. The Ez envelope is overlaid on the mixed-frequency square wave signal to better illustrate the phenomenon. Finite-element method computation of ot-TI using square-waves, in both A and B show the maximum envelope amplitude of Ez, located at the same point of the CA3 using the two different orientations, ML and AP. These two configurations are compared experimentally to determine the minimally invasive efficacy of ot-TI to generate beta/gamma range oscillations and SLEs in the mouse hippocampus

In the configuration shown here, we create electric fields parallel (medio-lateral, ML) and perpendicular (anterior-posterior, AP) to the axons in the mouse hippocampus. Additionally, as shown in Figure 1C and in supplementary Figure 2, it is not necessary to exclusively utilize sine waves for stimulation, a critical detail as clinical stimulation equipment most often uses square waves. The orientation-dependent configurations are shown with finite-element simulations of the mouse brain in Figure 1A and 1B, where the point of maximum envelope amplitude of the electric field (Ez component) is located at the same point of the CA3, but using the two different orientations, ML and AP.

## Materials and Methods

### Animals

All experiments were performed in accordance with European Council Directive EU2010/63, and French Ethics approval (Williamson, n. APAFIS 20359 - 2019041816357133). For this study we used 4 groups of 8 male OF1 mice (Charles Rivers Laboratories, France) aged 8-10 weeks. Animals were kept in transparent cages in groups of three to five, in a temperature-controlled room (20 ± 3°C) with a 12/12h night-day cycle. All animals had *ad libitum* access to food and water. Mice were divided into cohorts, differentiated by the electrode orientation: medio-lateral (ML) and anterior-posterior (AP) for both implanted and ot-TI. Additional mice were assigned for ot-TI studies with additionally implanted depth recording electrodes in the hippocampus to record the dynamics of the evoked SLE in the hippocampus vs the cortex. After the surgery, all mice were kept in separate cages to avoid fighting and to avoid damage to implanted electrodes.

### Surgical procedure

The 32 mice were anaesthetized via an intraperitoneal injection of ketamine (50mg/kg) and xylazine (20mg/kg) and placed in a stereotaxic frame with the head adjusted for bregma and lambda in the same horizontal plane. After midline scalp incisions, the following stereotaxic coordinates were used for craniotomies: [AP: −1.94, ML: +0.5; −0.5; −3.9; −4.3] for the ML electrode orientation and [AP: +2.2; +1.1; −4.4; −5.5; ML: −2.04]. For the implantable twisted-pair platinum electrodes (from PlasticsOne; wire length = 2mm, individual wire diameter = 125μm), coordinates were AP: −1.94, ML: 2.8, DV: 1.57 using a 15-degree angle. All the coordinates were calculated using the Paxinos Atlas. *Dura mater* was gently removed and four stainless steel mini-screws (Component Supply, Miniature Stainless Steel Self-Tapping Screws: TX00-2FH) were placed on the cortex without penetration into the brain tissue. Subsequently, dental cement (Phymep, SuperBond) was applied on the skull surface to fix the screws and the skull cap was formed using Dentalon. During the post-surgical recovery time (3 days), all the mice were observed for signs of pain, distress and neurological complications.

### TI Electrical stimulation

Electrical stimulation was performed using electrical stimulators (IntanTech, Intan 128ch Stimulation/Recording Controller) with two frequencies, 1200 Hz and 1250 Hz, and biphasic, bipolar pulses of 500 μs with the overall stimulation time of 10s (complete details in supplementary Figure 2 and 3). Additional details on isolation of stimulation systems is described in supplementary Figure 5. The intensity was gradually increased starting from 50 μA until seizures of stage 4 on the Racine scale were observed (ML) or until an animal exhibited intense motor response (i.e., convulsions in AP orientation). Control stimulations (1200/1200 Hz and 50/50 Hz) were performed at the threshold intensity of the TI session, with no observed SLEs. As expected, with no frequency offset, there is no envelope and thus no stimulation of the hippocampus. Each mouse received one full TI stimulation to identify the threshold and two subsequent control stimulations.

### Direct stimulation

After 3 days of recovery following implantation, a protocol to induce and to characterize the AD threshold was applied using the Racine scale. Following previous work, ADs were defined as high-amplitude spikes and polyspike epileptiform events visible after the applied stimulus. We used a 10 second stimulation, bipolar, biphasic, at 50 Hz with a 500μsec pulse width, with the amplitude increased in 50 μA steps, starting from 50 μA, until reaching the seizure of stage 4 on Racine scale. The animal was given 30 min rest between attempted stimulations.

### Behavioral evaluation

All mice were video monitored during stimulation. Behavioral responses were analyzed, taking care to distinguish between a motor response and an epileptic seizure. For seizure detection the following criteria were used, according to the Racine scale: Racine scale: Stage 0, no visible change in behavior; Stage 1, freezing with facial movements; Stage 2, head nodding; Stage 3, forelimb clonus; Stage 4, rearing without loss of balance; and Stage 5, rearing and loss of balance). The motor cortex response was evaluated, comparing the video recording with the simultaneous EEG recording, depth electrodes, and by extracting non-specific signs (limb twitching, vocalizing etc). In parallel to the LFP recording, video monitoring was continued during the complete stimulation/recording/rest sessions of the freely-moving animal in its cylinder open-field environment (50cm × 25cm × 25cm) to monitor the behavior and correlate it directly with the brain activity. Examples of Racine scale behavior after stimulation can be found on our Github (https://github.com/Florian139/Temporal-Interference.git).

### Virtual simulation

In order to have an idea of the stimulation in terms of applied voltage and electric field, COMSOL Multiphysics, version 5.5, (www.comsol.com) was used to create Finite-element method simulations of the electrodes and environment. Prior to simulations, we created using Hexagon software a fine 3D mouse brain based on the Paxinos mouse brain atlas. We embedded the stimulation electrodes (stimulation and grounds) as platinum stimulators in an environment having the frequency-dependent complex permittivity values of brain tissue. Using electrical stimulation from our electrodes in COMSOL with our mouse brain mesh, the equipotential surfaces and electric field lines applied by the electrodes could be visualized in the hippocampus. The orientation and positioning of the electrodes could be optimized using the simulation to better target the boarder of the CA3. The physical extent of field lines with respect to the orientation of the axons of pyramidal cells in the hippocampus could be better seen. The mesh geometry utilized and the basics on the simulation physics utilized can be found on our Github (https://github.com/Florian139/Temporal-Interference.git).

### Statistical analysis

All raw recordings were plotted and analyzed via Matlab. Statistical analyses (R software) of AD-thresholds were performed using the parametric one-way ANOVA and pairwise T-tests to calculate the probably significant differences in SLE amplitude, duration or severity between the 4 groups. For the TI group, the AD-threshold distribution appears to not follow a normal distribution. Therefore, all AD-thresholds were analyzed using a standard non-parametric Wilcoxon-Mann-Whitney tests. All necessary code to reproduce our figures, as lambda “.rhs” data file, can be found on our Github (https://github.com/Florian139/Temporal-Interference.git).

## Results

We induced SLEs in the mouse hippocampus using both TI and implantable electrode protocols. The target of stimulation was the border of CA3 and CA1 (detailed coordinates in the methods section), well-below the cortex surface.

As seen in Figure 2A-C for the two orientations of ot-TI, the maximum envelope is placed at the same location in the hippocampus, namely the border of the CA3 to CA1. Although the maximum is at the same point, the electric field lines are parallel (for ML) or perpendicular (for AP) to the axons of the hippocampus, the Schaffer Collaterals (SCs). In Figure 2D, for all ML mice, the shape of the evoked activity, the increase in the frequency range of beta/gamma (20-40 Hz), and the behaviorally observed seizures were all consistent with a classically-described stimulation-induced focal seizure of the hippocampus, however this is the first minimally invasive evoked SLE using the method of TI. All of the 8 ML mice exhibited a behavioral seizure 4 on Racine scale (overall threshold of 700μA per electrode pair) correlated with an electrophysiological SLE.

**Figure. 2:**
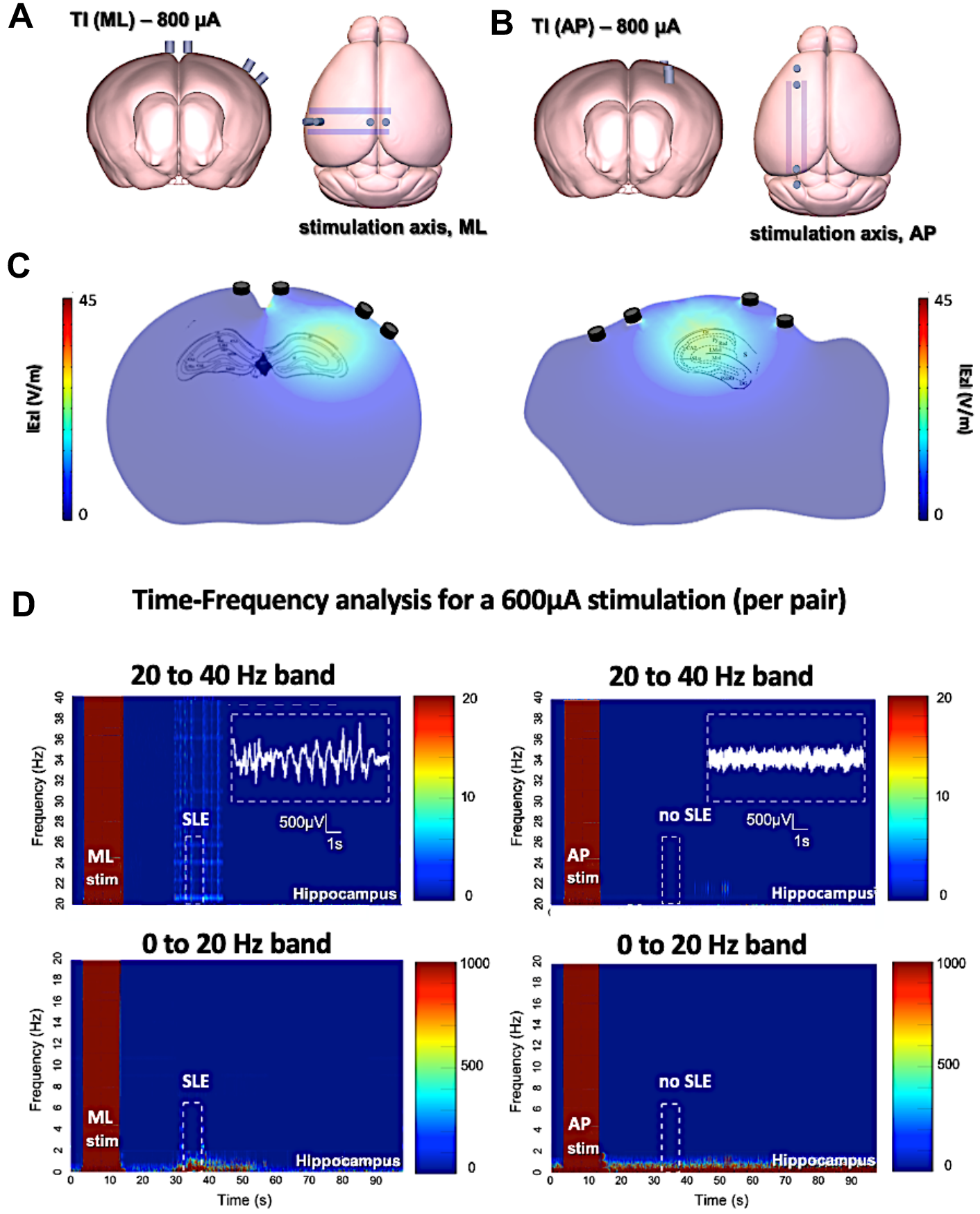
Evoked SLEs in the hippocampus of freely-moving mice. **(A)** and **(B)** depict the two ot-TI stimulation axes with electrodes to-scale. **(C)** The associated hotspots (maximum envelopes of electric field) are seen in the hippocampus, with the maximum amplitude placed at the CA3-CA1 junction as planned in the protocol. For ML/AP orientations, the electric field lines are parallel/perpendicular to the Schaffer collaterals (also see Figure 1). (**D**) Real biologically evoked SLEs in mice using ot-TI. For the ML orientation (left panels), parallel to the axons of the hippocampus, the shape of the evoked activity, the associated increase in frequencies of the beta/gamma range (20-40Hz), and observed behavioral seizures demonstrate a classic SLE, however it is the first minimally-invasive evoked SLE by using TI. The AP orientation (right panels), perpendicular to the axons of the hippocampus, showed no evoked SLEs, and for 8 mice only one showed a stage 4 behavioral seizure on the Racine scale with a threshold above 900μA. (additional details, video, and stimulation protocol in supplementary Figure S3 as in our Github). Cortex recordings shown in supplementary Figure 1.

In Figure 2D for AP mice, interesting things happen when the point of maximum envelope remains at the same coordinates, but the electric field is turned perpendicular to the axons of the hippocampus in a second group of mice. Clearly, no SLE activity is evoked with the AP orientation, although the amplitude of the envelope and its location are the same for both ML and AP. It is therefore necessary to increase the intensity of stimulation.

An explanation of stimulation using the AP orientation and its subsequent consequences in the motor cortex can be seen in Figure 3A-C, where the envelope of the electric field is compared to the underlying brain anatomy. In Figure 3A, the stimulation is equal to the stimulation applied in Figure 2 with no evoked SLE, and a maximum envelope of electric field is seen at the CA3-CA1 junction in the hippocampus with a small envelope in the neighboring cortex. In Figure 3B, the stimulation from the electrode pairs is increased and correspondingly the maximum envelope in the hippocampus increases, however so does the envelope in the neighboring cortex. This increase in stimulation results in Figure 3C, where a maximum limit of stimulation is reached as single-sided (corresponding to the implanted hemisphere) motor convulsions occur during the stimulation itself.

**Figure. 3:**
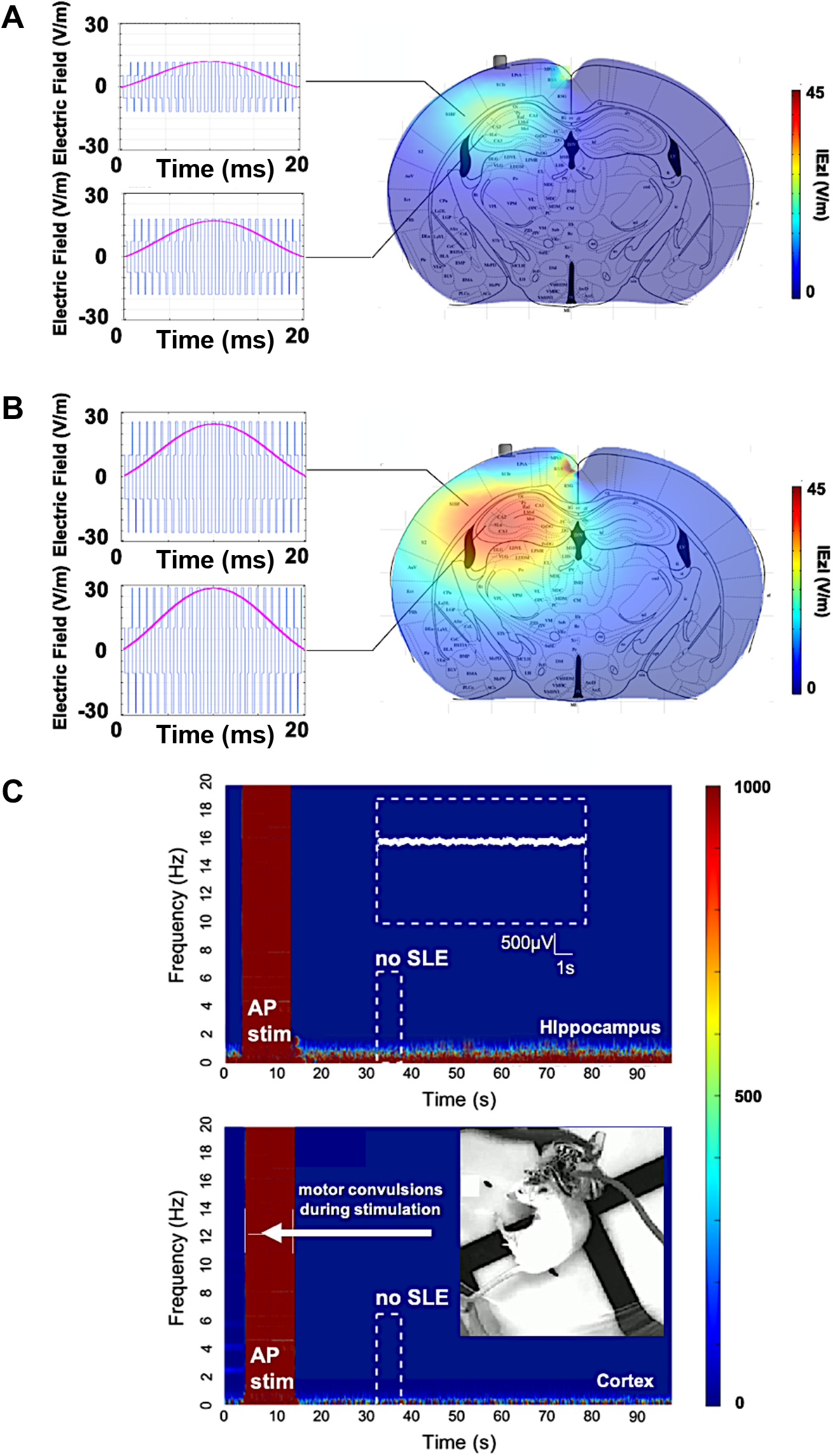
AP-ot-ti, non-preferential orientation, electric field envelope versus anatomy. **(A)** Profile of the electric field and envelope for the stimulation in Figure 2B (600μA per pair), and **(B)** stimulation with the same orientation but with a higher stimulation amplitude (1500μA per pair). Clearly, continuously increasing the intensity of stimulation will increase the envelope of stimulation in the hippocampus, ideally leading to an SLE. However, the increase in stimulation amplitude creates a non-trivial increase of the envelope located in the cortex. **(C)** This envelope in the cortex is enough to evoke arbitrary activity and motor convulsions. As the stimulation intensity is increased for the AP orientation, significantly past the threshold for SLE in part B, single-sided motor convulsions are observed during the stimulation itself.

In summary, in Figure 3A, corresponding to the stimulation applied in Figure 2, the amplitude of the electric field is clearly higher at the focal point of the coordinates in the hippocampus. However, due to the non-preferential orientation of the electric field to the axons of the hippocampus, the threshold to evoke an SLE has not been reached. As amplitude increases to find the threshold for an SLE, a limitation of the TI method is observed with direct consequences for uncontrolled or indiscriminate orientations of TI electric fields.

Namely, as the amplitude of the focal spot increases for non-preferential orientations, so does the envelope along a radial axis moving away from the focal spot. A target structure along the radial path may have a threshold for activation, in this case convulsions in the motor cortex, which will be reached before the threshold for activation of the target, in this case seizure activity in the hippocampus. This understanding can be algorithmically formulated and used to tailor stimulation orientations, not limited to AP/ML, for other subcortical structures.

To support the existence of orientation-dependent thresholds in TI, we subsequently demonstrate the dependence of orientation on thresholds using implantable electrodes. As seen in Figure 4A and 4B, we see that when using classic implanted stimulation electrodes, the threshold to evoke SLEs is highly dependent on the orientation of the electric field. For a preferential ML orientation with the equipotential surfaces of the electric field parallel to axons (where a gradient of potential is created along the axons), the threshold requires half the injected charge. For a non-preferential AP orientation with the equipotential surfaces of the electric field perpendicular to axons (where no gradient of potential is created, axons are fixed at one potential surface), the threshold requires double the injected charge.

**Figure. 4:**
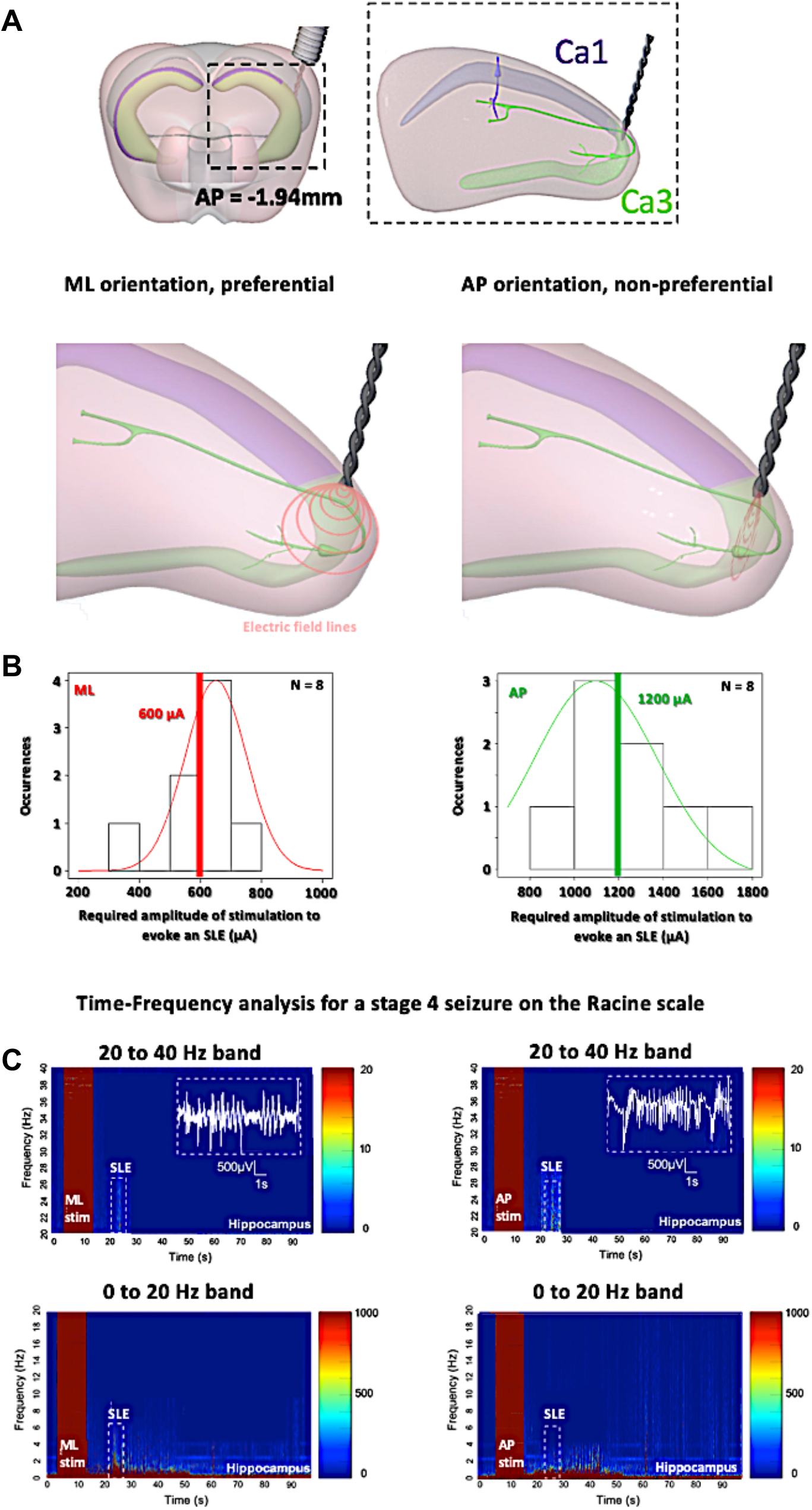
Orientation-controlled stimulation of SLEs in the hippocampus with classic implanted stimulators. **(A)** Stimulation at the boarder of CA3 and CA1, for implantable electrodes with preferential ML orientations having equipotential surfaces of electric field parallel to axons, and with non-preferential AP orientations having equipotential surfaces of electric field perpendicular to axons. **(B)** The threshold to evoke SLEs with ML requires half the injected charge, compared to AP. This is consistent with our TI findings, that the AP orientation is more difficult for evoking SLEs **(C)** As soon as the stage 4 on the Racine scale is reached, an increase in power intensity is shown with a peak in beta/gamma range oscillations between 20-40 Hz, correlating with SLEs (15s after the stimulation) for implantable orientations, however for AP (right panels) double the current is required.

To be sure that we induced the same SLEs in the two implantable groups, we analyzed and compared the electrophysiology and the behavior (Figure 4C). All mice from the two implantable groups, both ML and AP, exhibit electrophysiological SLEs, with an increase in the beta/gamma band (20-40 Hz), and stage 4 seizures on the Racine scale, however with a threshold which is significantly lower for the ML orientation. This is an interesting result, as no motor convulsions are observed for the AP orientation with implantable electrodes because there is no axis of stimulation through the cortex, allowing additional insight into the difference in thresholds for preferential and non-preferential orientations of stimulation. Correspondingly, we analyzed and compared the electrophysiology and the behavior for the implantable ML orientation and ML-OT-TI orientation (Figure 5).

**Figure. 5:**
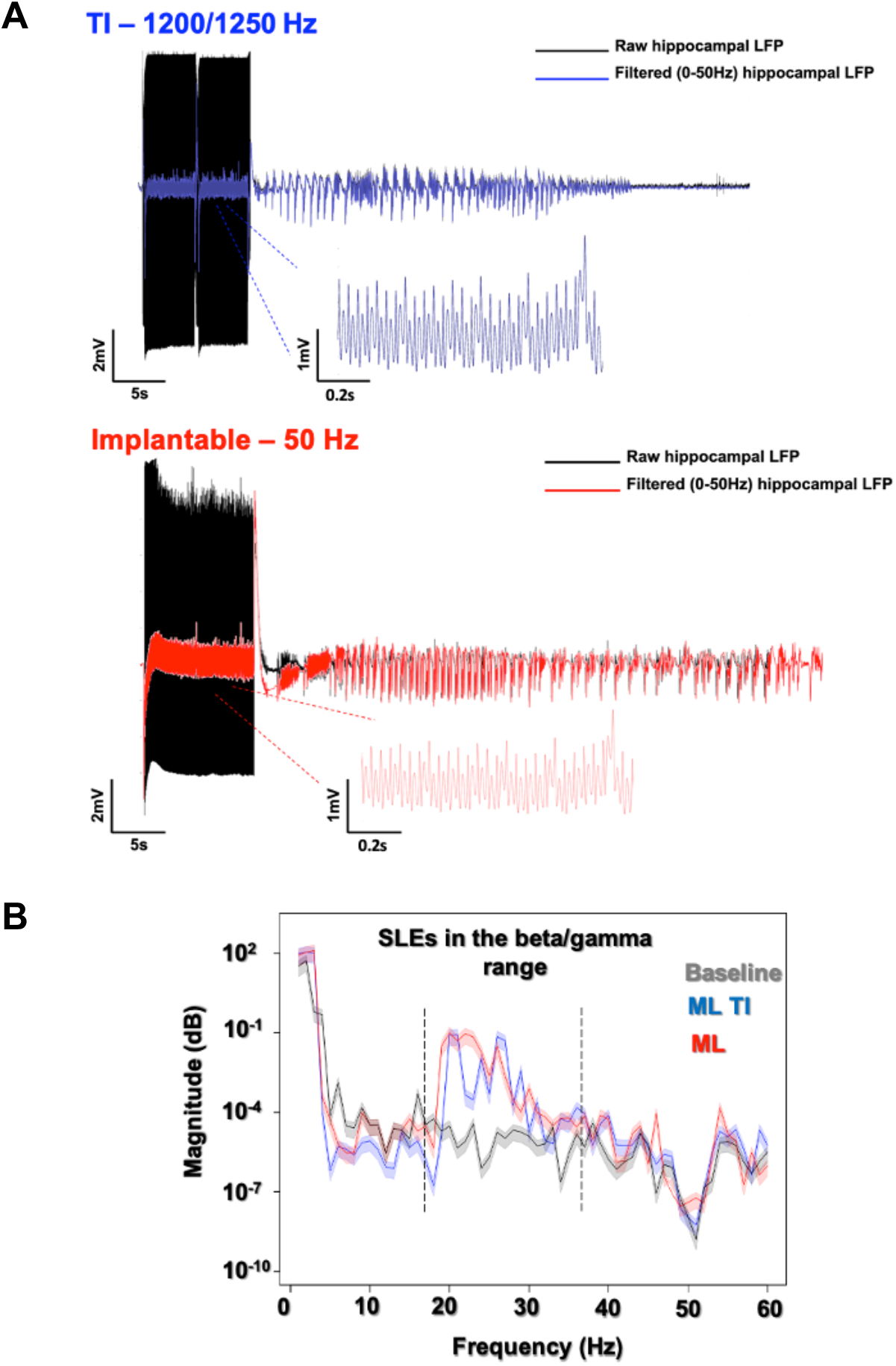
LFP recording of freely-moving mice after a 600μA stimulation using the preferential ML orientation. All recordings (ML-ot-Ti in blue and ML-implantable in red) were analyzed using a 50Hz lowpass filter to clearly distinguish the SLE. **(A)** The classic implantable stimulator provides a 50Hz applied stimulation in the hippocampus. TI provides a 50Hz stimulation, corresponding to the differences between the two stimulation frequencies in the hippocampus (1200 and 1250Hz). **(B)** Power Spectral Density (PSD) shows SLEs (15s after the stimulation) with a peak in the 20-40Hz band (Beta/Gamma) characteristic of a rapid discharge for both implantable and TI stimulation protocols.

In Figure 5, electrophysiological SLEs and the associated power spectral density (PSD) are shown, where the PSD is 15s after the stimulation and seen to have a peak power in beta/gamma range between 20 Hz and 40 Hz, true for implantable ML and identical for the ML-ot-Ti. In Figure 5 the amplitude of the evoked SLEs for both implantable orientations is statistically the same with the ML-ot-Ti (friedman-test, p-value > 5%).

The 3 groups with detectable SLEs (AP implantable, ML implantable, and ML-ot-Ti) are further compared using the amplitude, duration, and severities of these SLEs. For all SLEs detected, we measure the amplitude and the duration of the SLE, as seen Figure 6. We compared amplitudes and duration of 20 SLEs per group. Figure 6A shows the SLE amplitude distribution for the 3 groups. Using first a normality test (Shapiro) and then a T-test, there were no differences between the 3 groups. SLEs durations show no differences between ML implantable and ML-ot-Ti, however these 2 groups are statistically different from the SLE durations of the AP implantable group. Consistent with the previous result, the severity of the seizure is also lower on all Racine scale stages for the AP implantable. It is simply more difficult to induce seizures with an AP orientation, and the induced seizures are also shorter and less severe than the seizures with ML orientations. However, the two ML groups, implantable and TI, are statistically identical in term of induced seizures, amplitudes, durations, and severities meaning that the minimally-invasive method is generating the same electrophysiological and behavioral event, but without the necessity of implanting the tissue.

**Figure. 6:**
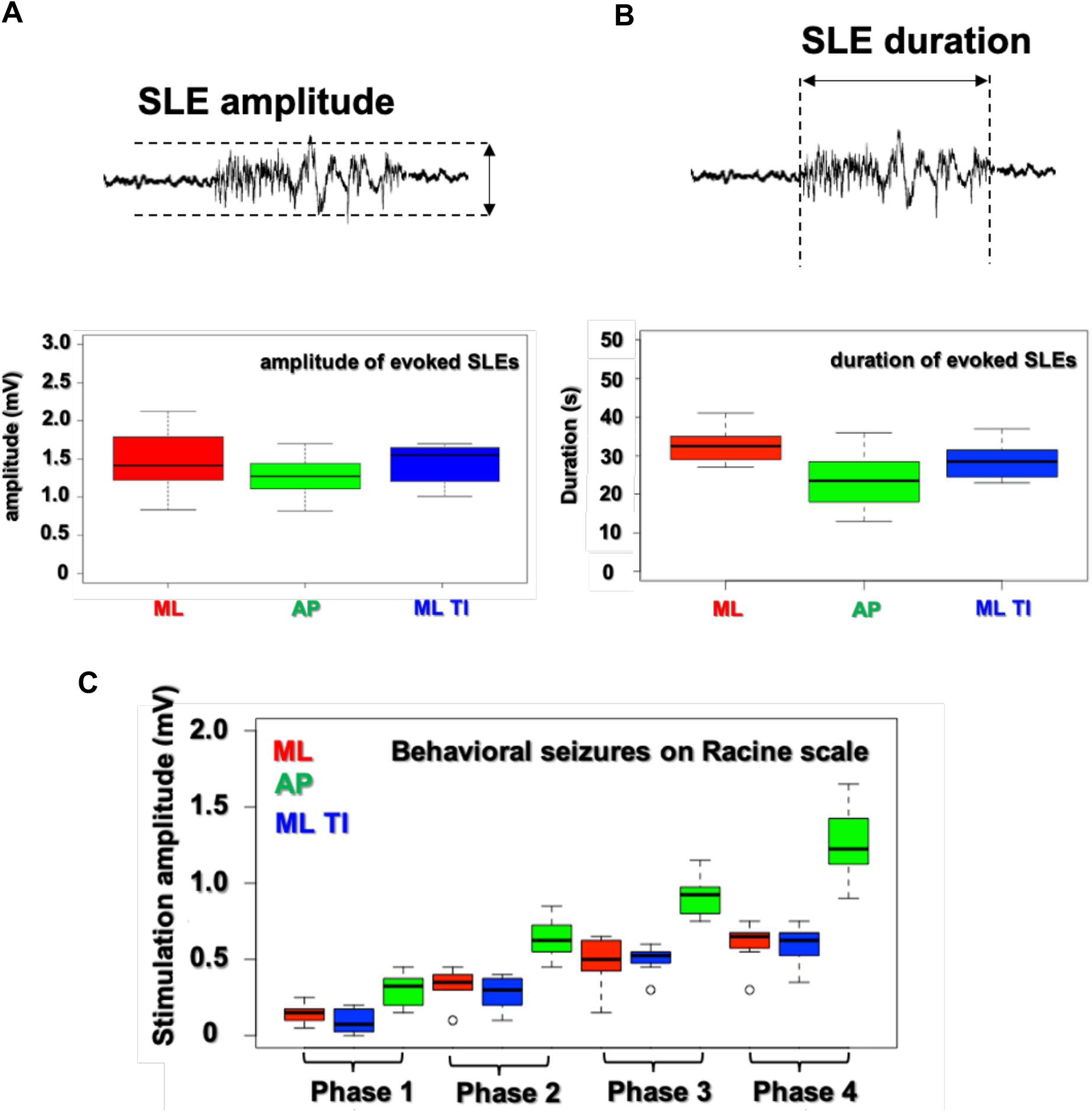
Analysis of SLE characteristics (amplitude, duration, severity) in the hippocampus. All SLE detected have been characterize in amplitude, duration and severity. It appears no differences between ML implantable and ML-ot-Ti (p-value > 5%) for SLE amplitudes, durations and severities. However, we could find some differences in duration and behavioral output for the AP-implantable orientation. Seizures are more difficult to induce with the AP orientation and the AP-induced seizures are shorter in time.

Indeed, orientation dependence in TI is not surprising, as orientation dependent stimulation is known to exist for implantable electrodes in the field of Parkinson’s disease^12,13^. The main point is that identical evoked SLEs in the beta/gamma range can be attained in the hippocampus using temporally interfering electric fields, without the necessity of an implanted stimulator, if care is taken to account for the orientation of the electric field with respect to the axons of the hippocampus. We created both orientations of preferential ML and non-preferential AP mice using ot-TI. Based on our calculation of geometry and field strength, we can locate the focus at the exact intersection of the CA3 and CA1 used in the implanted stimulation experiment.

## Discussion

Clinically, comparing electrical activity from different brain regions, in particular electrically stimulated brain regions, in real-time as seizures emerge is fundamental to the identification of the EZ^14^. Seizure onset is characterized by dramatic changes in brain rhythms with several patterns of onset often observed, namely preictal epileptic spikes, trains of spikes, rapid-discharges, and slow-wave complexes with frequencies involved in the range of beta/gamma (15–30 Hz, 20-40 Hz in mesial temporal seizures)^15,16^. Typically, 8–15 implants are located in patient’s brain (generally 0.8 mm diameter, with multiple electrode contacts 2 mm long and 1.5 mm apart), allowing stimulation at various mesial and lateral sites usually using lower frequencies (typically 1–50 Hz, for 5s) to map functional cortex or to evoke seizure onset^17^. Unfortunately, there are some parts of the brain which remain unreachable for electrode implant due to the presence of blood vessels, eloquent cortex, or other surgically-complicated implant locations and possible subsequent deficits. Clearly, there are limitations to the number and placement of intracranial electrodes and, as we have demonstrated here, ot-TI could be an interesting and additional tool for evoking and stimulating seizure onset. The possibility to explore some parts of the brain unreachable by SEEG electrodes could be very interesting during clinical investigation of, for example, the insula as it has a high density of arteries and veins and it is well-known to be a very difficult tissue area of the brain to operate^18^. This is important because the insula is also well-known to be implicated in refractory partial epilepsies and can be part of the epileptogenic zone of the patient^19,20^. Unfortunately, it is commonly described that patients cannot undergo a proper functional mapping of this site because of the risk of hemorrhage during implant^21^. Correspondingly, ot-TI also has the possibility to reduce the overall number of SEEG electrodes implanted. There are numerous well-documented complications in SEEG electrode surgery, related infections, and complications which can results in necessary explanation of the electrode^22,23^. A reduction in SEEG electrodes would correspondingly reduce the number of SEEG-related complications.

In summary, TI is a very recently introduced concept for brain stimulation^11^. In the work here, we have utilized a new orientation-tunable form of TI (ot-TI) to evoke SLEs in the mouse hippocampus having the range of beta/gamma frequencies seen in MTLE, but without the necessity of implantable electrodes. Our target was the classic CA3-CA1 border structure used in epilepsy studies where the neurons of the CA and their axons, the Schaffer collaterals, are a well-known target for evoking seizures in vivo and in vitro with penetrating electrodes^24,25,26^. In our alternative technique relying exclusively on nonpenetrating surface electrodes and ot-TI electric fields, we have shown that the SLEs generated are indeed focally evoked in the hippocampus and identical to focally evoked SLEs generated with classically implanted electrodes. We additionally demonstrated the use of square-waves with TI stimulation, as clinically, square-waves are most often used in stimulation. As we have shown, the orientation of the electric field plays a key role in evoking events, and if carefully controlled, optimal orientation dramatically lowers thresholds for both implantable electrodes and correspondingly the minimally-invasive ot-TI, replicating the implantable stimulation without the need of penetrating the cortex. This method has promise to significantly advance our capacity of probing the organization of spatiotemporal brain activity and could dramatically increase the explorable tissue for clinical definition of the EZ. Finally, it could make it possible to deliver therapeutic stimulation in a highly controlled and accurate non-invasively to deeper regions of the brain, an interesting topic for a subsequent study. Regardless, the technique undoubtedly holds great promises with potential applications in epilepsy, but also for a wide range of other brain disorders currently managed by electrical brain stimulation.

## Acknowledgments

A.W. acknowledges funding from the European Research Council (ERC) under the European Union’s Horizon 2020 research and innovation program (grant agreement No 716867). A.W. and E.R. acknowledge funding from the Excellence Initiative of Aix- Marseille University—A*MIDEX, a French “Investissements d’Avenir” program. A.W. acknowledges support from the Knut and Alice Wallenberg Foundation.

## Author Contributions

A.W. conceived the project. E.R., E.A., and F.M. performed experiments. B.B. performed finite-element simulations. F.M. analyzed neural data. A.W. wrote the paper with input from the other authors, including R.C., V.J., A.T., and F.B.

## Supplementary Figures

**Supplementary Figure S1:**
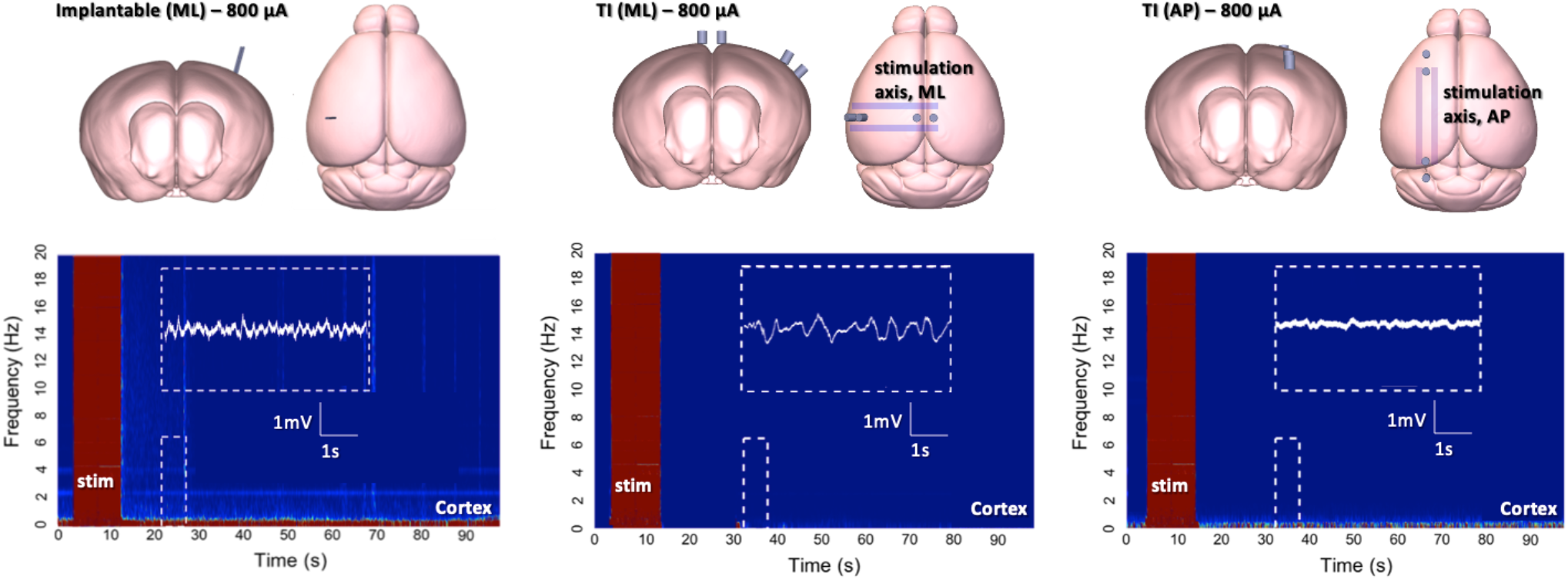
No induced SLE in the cortex: In all our experiment and as a control of no cortical stimulation, we recorded from a recording electrode directly on the cortex. In all cases, even if an SLE was induced in the hippocampus, the cortex was never activated (for amplitudes < 1500μA).

**Supplementary Figure S2:**
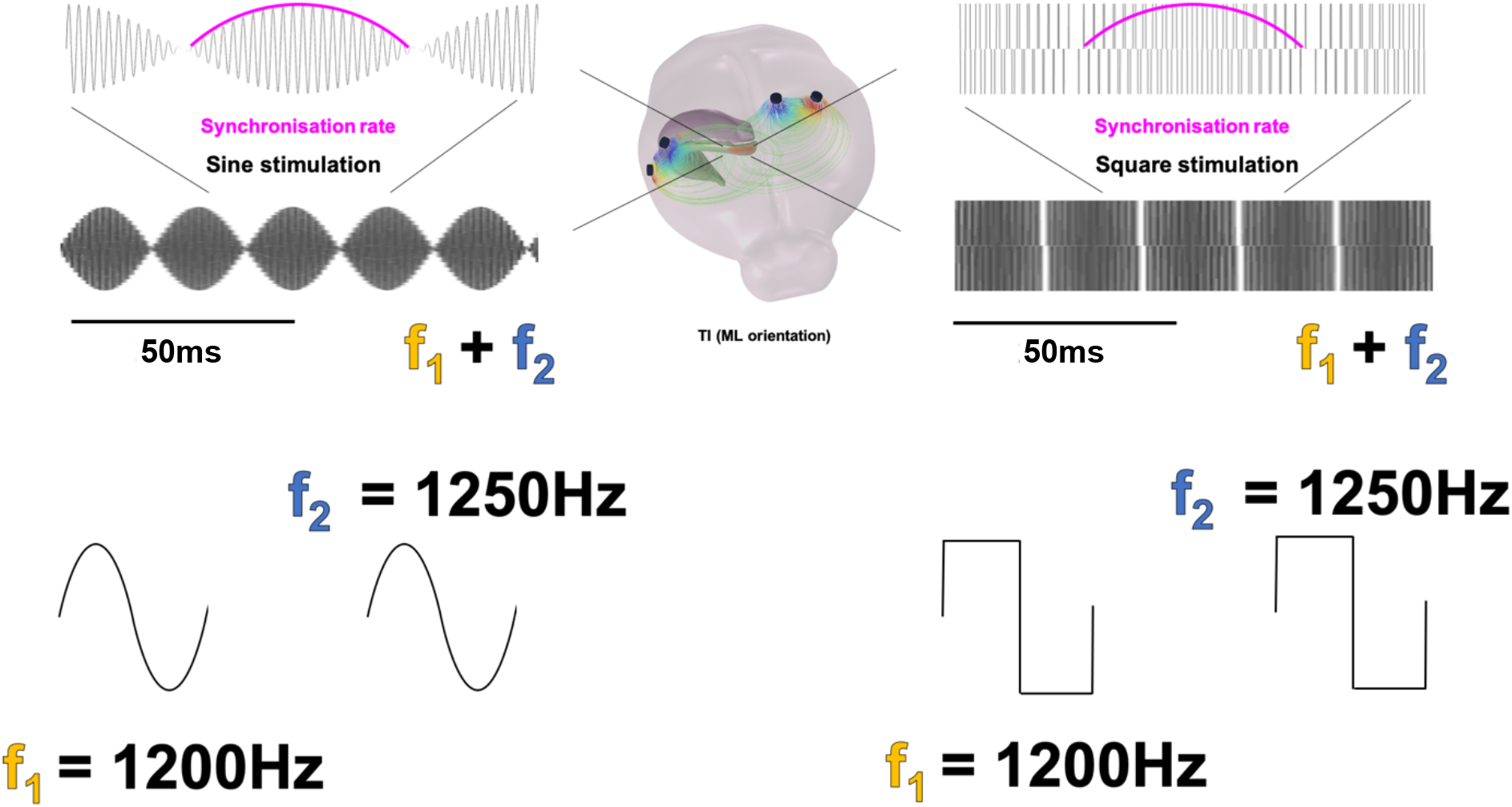
Square-wave TI. Using square waves for TI is the same as using sine waves. Practically, this does not appear as an envelope on the screen during stimulation, rather it appears as a “synchronization rate” – essentially the two square pulses coming closer and closer into phase. However, when the signal is filtered, one will visualize the low frequency envelope as expected.

**Supplementary Figure S3:**
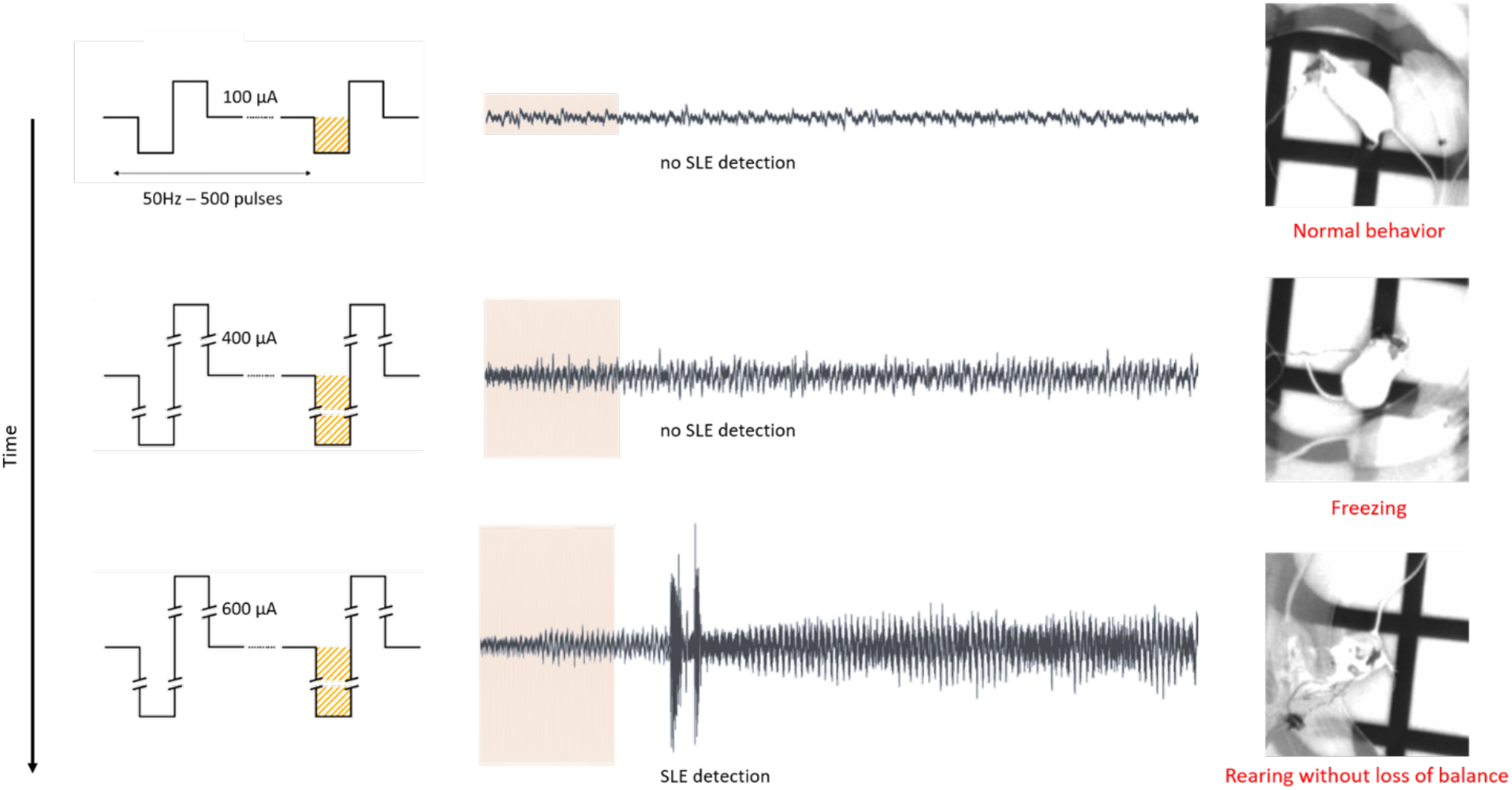
Stimulation Protocol for TI and implantable. Each session consists of the 10s stimulation and 5min EEG/video monitoring, during which animal’s behavior is observed and analyzed, in accordance with simultaneous EEG-recording. Once an SLE is detected, the stimulation threshold is noted, no further stimulation is necessary.

**Supplementary Figure S4:**
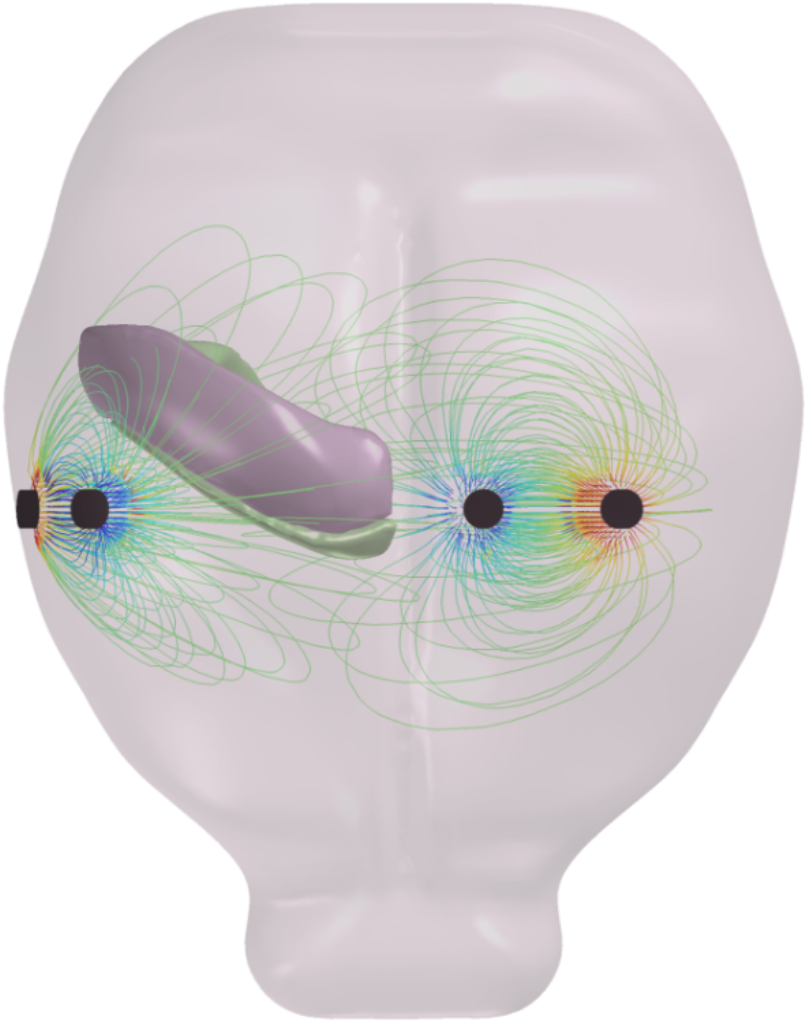
Field lines of TI. The field lines in Figure 1 of the main manuscript are plotted to highlight the symmetry of the TI field with respect to the orientation of axons in the CA of the hippocampus. Complete plots of the field lines, as show here for the ML orientation, also include lines at angles deviating from the primary symmetry. However, as pictured in Figure 2 of the main manuscript, the field lines along the axis of symmetry have the largest relevant envelope due to the orientation of the two pairs of electrodes.

**Supplementary Figure S5:**
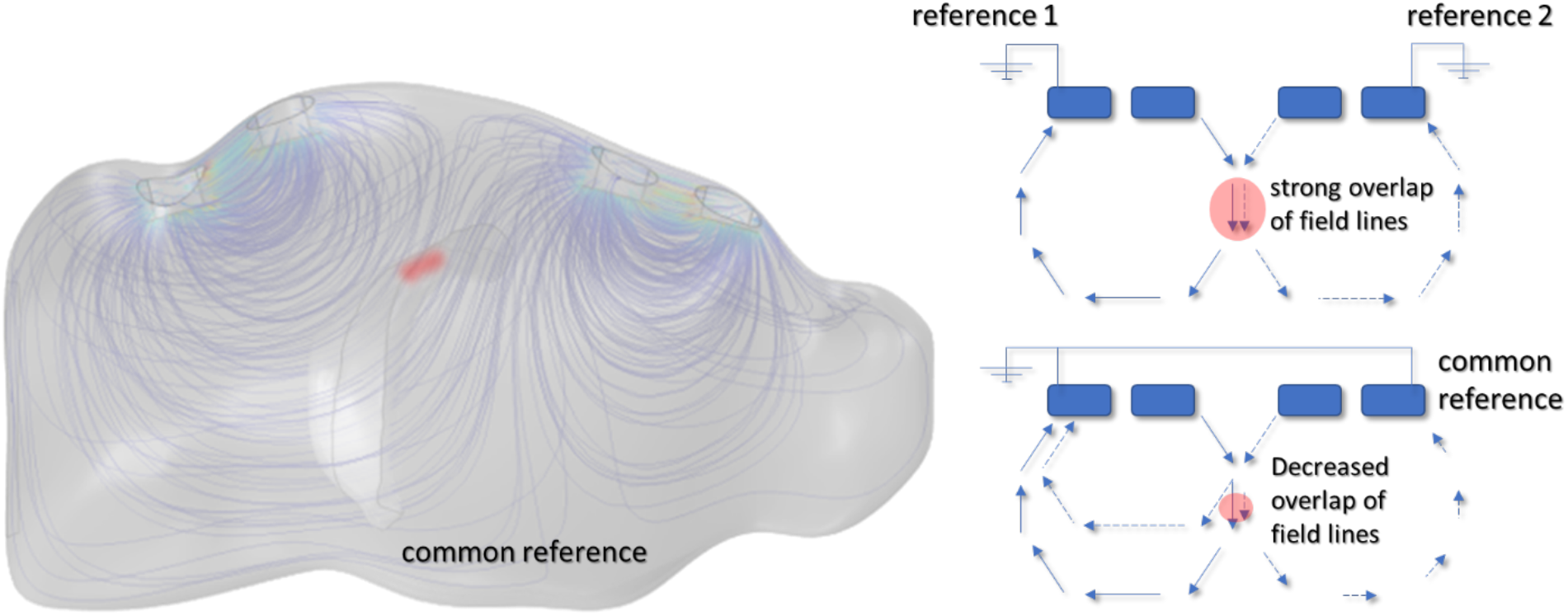
How NOT to make a TI field. It is extremely important that the two stimulation systems are isolated. If the stimulation systems share the same reference/ground the situation pictured in the figure here will occur. Field lines from the cathode of one stimulation pair will travel to the anode of the opposite stimulation pair. This will significantly weaken, and possibly completely remove, the point of the maximum envelope of stimulation (red). This can often not simply be done by connecting two separate stimulators, as both machines will be connected to the same wall socket. Ideally two separate battery-powered stimulators will provide the perfect isolation.

